# Comparative analysis of the structure of complete chloroplast genomes in genus Mangifera and accuracy verification about phylogenetic analysis based on gene *ycf*2 in genus level

**DOI:** 10.1101/2022.04.05.487216

**Authors:** Yujuan Tang, Shixing Luo, Yu Zhang, Ying Zhao, Riwang Li, Limei Guo, Guodi Huang, Aiping Gao, Jianfeng Huang

## Abstract

Mango is an evergreen plant belonging to the genus *Mangifera* of the Anacardiaceae family. Genus *Mangifera* has 69 species of *Mangifera* around the world that are mainly distributed in tropical and subtropical countries, including India, Indonesia, the Malay Peninsula, Thailand, and South China. It is a popular tropical fruit known as the “King of Tropical Fruits”. However, the study of the structure information of the complete chloroplast genome of *Mangifera* was microscopic, there was no report about the comparison of SSR, Ka/Ks, codons analysis and RNA editing, so in this study, we sequenced the 6 *Mangifera* samples and used three different ways to analyze the relationship of 6 species of *Mangifera*. Then we got some results, through the RNA editing and Ka/Ks calculating, we found the species could be divided into two groups, and the difference between the two groups was protein-coding gene *ccsA*. Moreover, all RNA editing occurred conversion of C to T and the gene *ndhB* had the most RNA editing sites in all species. In Ka/Ks analysis, the gene *atp*B, *cem*A, *clp*P, *ndh*D, *pet*D, *pet*B and *ycf*15 would be suffered from the positive selection after divergence. We also find the IR regions in these seven samples were very conservation through IR contraction and expansion and Sequence Divergence Analysis. Finally, we tried to confirm the relationship between 7 samples of *Mangifera* in Angiosperms in 3 different ways. Then we got that ML210 and MP090 had a closer relationship than others, MS796 had a closer relationship with ML210 and MP090 than others. At the same time, the method of phylogenetic analysis based on the gene *ycf*2 was not more accurate at the genus level than the method based on complete cp genome and proteincoding genes.

## 1 Introduction

Mango is an evergreen plant belonging to the genus *Mangifera* of the Anacardiaceae family. Genus *Mangifera* has 69 species of *Mangifera* around the world that are mainly distributed in tropical and subtropical countries, including India, Indonesia, the Malay Peninsula, Thailand, and South China [1,2]. Its leaves have two colors: the fuchsia color for new leaves and the green color for old leaves, and it is a popular tropical fruit [3,4]. And it was widely planted because of its high economic and nutrient value [5,6]. Because of its unique taste and pleasing appearance, it is widely loved by fruit lovers and is known as the “King of Tropical Fruits” [7]. Southeast Asian countries have a very long history of mango cultivation that spans thousands of years [8]. Mango spread to Africa, South America hundreds of years ago, and now they have developed several cultivated species suitable for their local climate [9,10].

As active metabolic centers responsible for photosynthesis and the synthesis of amino acids, nucleotides, fatty acids, phytohormones, vitamins, and other metabolites, chloroplasts play essential roles in the physiology and development of land plants and algae[11,12]. In most land plants, the chloroplast genomes exhibit highly conserved structures and organization and typically exist as circular DNA molecules with a size of 120–170 kb [13]. Chloroplast genomes generally have a quadripartite structure and contain a large single copy (LSC) region and a small single-copy (SSC) region separated by inverted repeats (IRs). However, these IR regions are missing in some species[14,15]. Specific characteristics of the chloroplast genome, such as its maternal inheritance, haploid nature, and low level of recombination, make it a robust tool for genomics and phylogenetic studies of several plant families[16–19]. Moreover, considerable variations within chloroplast genomes can also provide helpful information for evaluating the phylogenetic relationships of taxonomically unresolved plant taxa and understanding the relationship between plant nuclear, chloroplast, and mitochondrial genomes in plants[18,20–22].

Phylogenetic analysis of *Mangifera* species has been a hot topic of research [23,24]. At the same time, the complete chloroplast genome sequences can provide more genetic information and higher species resolution ability than other molecular data. However, the study of the structure information of the complete chloroplast genome of *Mangifera* was very little, and there were no information about SSR, Ka/Ks, codons analysis and RNA editing. In this study, we sequenced six samples of *Mangifera* and analyzed them from codon usage, selection pressure, RNA editing, Repeat Sequences and SSR, Polymorphism Analysis, Comparison of Genome Structure and Phylogenetic analysis base three different ways.

## 2 Materials and Method

### 2.1 Plant Material

There are six samples in the genus *Mangifera* in this study. In these six samples, 3 of them were collected from Guangxi, China (*Mangifera indica-OK104092* was collected in Hezhou, *Mangifera persiciforma-OK104090* in Baise and *Mangifera sylvatica* OK104091 in Nanning), one species was collected in Yunnan, China (*Mangifera Siamensis*-MZ926796 in Xishuangbanna); 1 species was from Thailand (*Mangifera indica*-OK104089 in Chiengmai) and another was from U.S.A. (*Mangifera odorata* was in Florida).

### 2.2 DNA Extraction, Sequencing and Annotation

Fresh leaves of the plants were collected and immediately stored at −80 °C. Total genomic DNA was extracted using the modified CTAB method [25]. The integrity, quality, and concentration of the DNA were determined by agarose gel electrophoresis and a NanoDrop spectrophotometer 2000 (Thermo Fisher Scientific, Waltham, MA, USA). Then high-quality genomic DNA was used to construct libraries with an average length of 350 bp using the NexteraXT DNA Library Preparation Kit (Illumina, San Diego, CA, USA) and sequenced on the Illumina Noveseq 6000 platform (Illumina). More than 11.4 million paired-end reads of each sample with an average length of 150 bp were generated and edited using the NGS QC Tool Kit v2.3.3[26]. The complete circular assembly graph was checked and further extracted by visualization of the GFA graph files assembled from SPAdes 3.11.0 software [27]. These chloroplast sequences were annotated using PGA[28], and BLAST was used to evaluate the results.

### 2.3 codon usage, selection pressure and RNA editing analysis

We selected all the PCGs in 7 samples using Unipro ugenes v.36.0[29]. Then codon usage frequency in each of the seven samples was analyzed using codonW v. 1.4.2 (http://codonw.sourceforge.net/)[30]. Synonymous codon usage and relative synonymous codon usage (RSCU) were conducted to determine if the plastid genes were under selection. Then we aligned the PCGs by blsastn with setting evalue: 1e-5, max_target_seqs: 1, outfmt: 6 and converted the files to PAML format. PAML was used to analyze Ka/Ks with the default setting [31]. Finally, we used PREP software[32] to identify the RNA editing sites by comparing the PCGs genes in 7 samples and the cut-off score was set as C=0.8 to find all actual RNA editing sites.

### 2.4 Identification of Repeat Sequences and Simple Sequence Repeats (SSR)

SSR identification was detected on the chloroplast genome sequences by MIcroSAtellite identification tool with the parameter settings: unit-size (nucleotide) _min-repeats: 1_8, 2_5, 3_4, 4_3, 5_3, 6_3. The minimum distance between two SSRs was set to 100 bp [33]. Dispersed (forward, reverse, palindrome, and complementary) repeats were determined by running the REPuter program with a minimum repeat size of 30 bp and similarities of 90%. Tandem repeats were identified by running the web-based Tandem Repeats Finder, with alignment parameters set to 2, 7, and 7 for matches, mismatches, and indels, respectively[34].

### 2.5 Polymorphism Analysis, Comparison of Genome Structure, and IR Region Contraction

The chloroplast genome sequences were aligned by MAFFT v7.4.29 [35] with MI098 as references and compared with the chloroplast genomes of others using the mVISTA program in the Shuffle-LAGAN mode [36]. A sliding window analysis was conducted for nucleotide variability (Pi) in the complete chloroplast genome to identify the rapidly evolving molecular markers using DnaSp v6.12.03 [37]. The window length was set to 800 bp, with a 100 bp step size. Then we use IRscope to analyze IR Region Contraction. (https://irscope.shinyapps.io/irapp/)

### 2.6 Phylogenetic Analysis of Chloroplast Genomes

We used three ways to construct a phylogenetic tree. We used 25 complete chloroplast genomes to build the phylogenetic tree in the first one. We used the gene *ycf*2 in 25 species to construct the phylogenetic tree in the second way. Thirdly, we just used 86 PCGs of 7 *Mangifera* species. Firstly, we downloaded 18 complete chloroplast genomes from NCBI and we selected the gene *ycf*2 from all 25 chloroplast genomes and 86 PCGs of 7 *Mangifera* species by using Unipro ugenes v.36.0[29]. Secondly, we separately aligned gene *ycf2* selected from 25 species, 86 PCGs from 7 chloroplast genomes of *Mangifera* and the 25 complete chloroplast genomes (18 downloaded chloroplast genomes and seven samples in study). In this step, the software we used was Mafft7.4.29 and the strategy was FFT-NS-2. Finally, we used the model finder to select the TVM+F+I+G4 model[38] and constructed the phylogenomic tree by IQtree 2.0[39] with 1000 bootstrap and maximum-likehood method. During building the phylogenomic tree, we used *Citrus aurantiifolia* (KJ865401) as an outgroup in the final step. The first two ways used the same software and setting; however, we did not use any outgroup in the third way.

## 3 Result and discussion

### 3.1 Genomic Characteristics of Chloroplast

First of all, we will use the following shorthand instead of the full name: *Mangifera* indica-OK104089 (MI089), *Mangifera* indica-OK104092 (MI092), *Mangifera* Siamensis-MZ926796 (MS796), *Mangifera* odorata-MZ926795 (MO795), *Mangifera* persiciforma-OK104090(MP090), *Mangifera* sylvatica-OK104091(MS091) and *Mangifera* longipes-MN917210 (ML210). The size of the complete chloroplast of 7 samples was around 157,604 bp to 158,889 bp. The largest one was MO795, the smallest one was MS796 and the difference was 1,285 bp. After comparing seven chloroplast genomes, we found two interesting results. Firstly, the size of the inverted repeat region (IR region) in 7 samples was very similar, and they were around 26360 bp to 26389 bp. MI089, MI092, M0795 and ML210 had the same size IR regions. It meant that the difference in chloroplast size did not depend on the IR region. We also found that the difference of large single copy size in 7 samples was around 1201, so it showed that the size of LSC could affect the whole chloroplast genome size. Secondly, the GC content of 7 chloroplast genomes covered around 37.82% to 37.9%, and they increased as the size of chloroplast genomes decreased (Table 1).

**Table 1.**
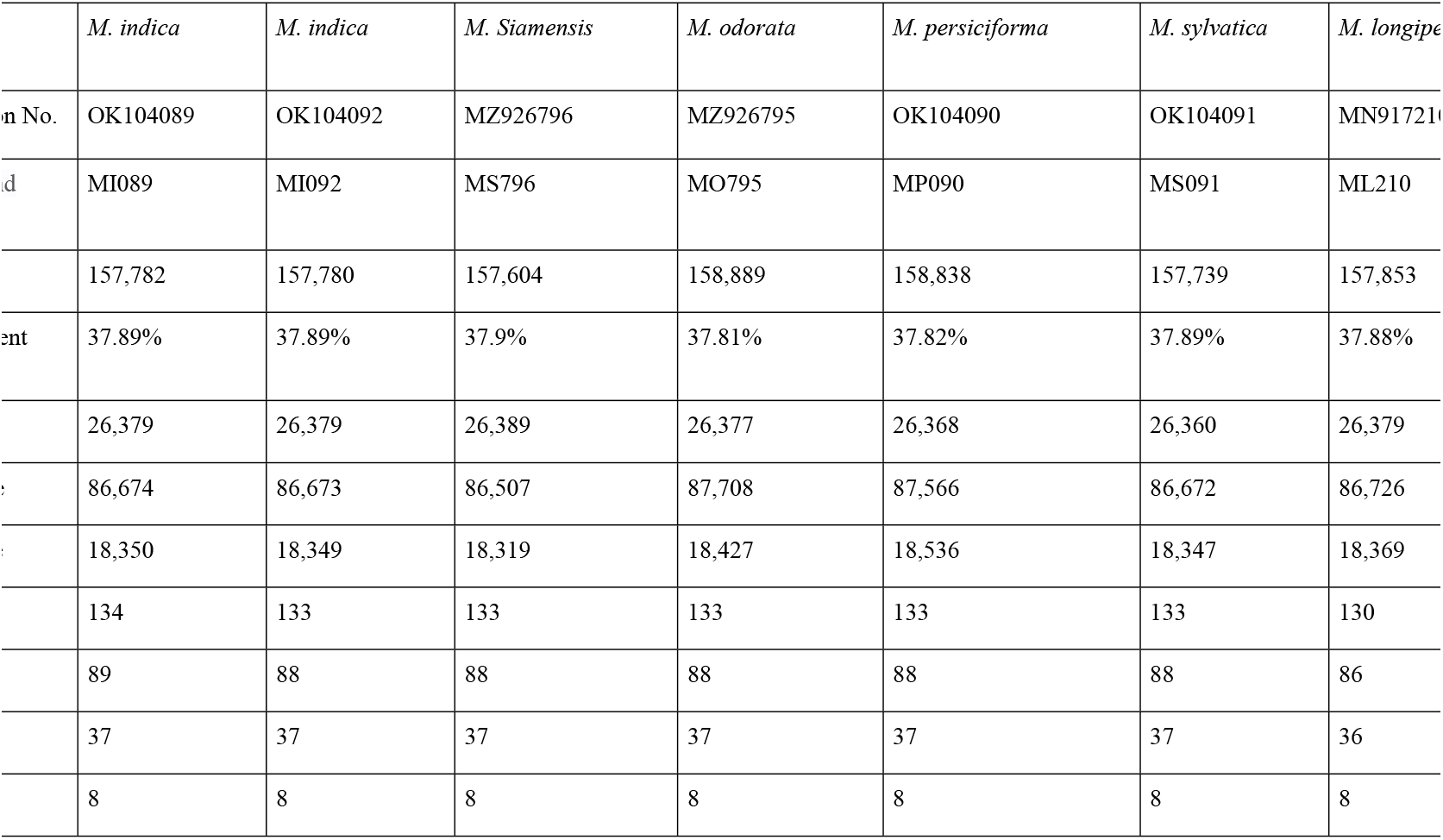
The feature of 7 complete chloroplast genomes.

We also detected around 133 to 134 genes in 6 species; they all had 37 tRNA. In addition, MI089 had 89 protein-coding genes and others had 88 protein-coding genes, so the total genes of MI089 were 134 and others were 133 (Table 1). All species in this study had eight rRNA in their chloroplast genomes.

### 3.2 Codon Usage and RNA editing analysis

In this study, we found some interesting things that the number of codons was independent of the size of the chloroplast genome. The largest chloroplast genome was 158,889 bp and from MO795, but it did not contain the most codons. However, it had the second most codons (26594 codons) or 79782 bp total protein-coding genes. And the chloroplast genome of MP090 contained the most codons, and the quantity was 26611 codons. However, the size of this chloroplast genome was 158,838 bp. We also found that the smallest chloroplast genome did not contain the least codons. It was MS796 containing 26578 codons, more than species ML210 and MS091. It meant that the size of the chloroplast could not state the size of total PCGs in chloroplast genomes in the same genus. The number of codons from the species in the study was between 26508 and 26611, and this meant that the range of the total size of protein-coding genes would be between 79524 and 79833 bp (Table S1).

Then we analyzed the codon usage in all seven samples and found that there were very similar the percentage of amino acids in different species in this study, Leucine was the most amino acid, around 10.56-10.59% in all of the species in this study, and it covered about 2,798 to 2,804 codons. Cysteine was the most minor amino acid in the species, around 1.17-1.18% and covered 310 to 314 codons. (Figure 1). Furthermore, we also found the difference in the size of PCGs in these seven samples was around 309 bp. Usually, the species in the same genus could not have a significant difference in the size of the PCGs, even the size of the chloroplast genome. However, the difference between the largest and smallest size of PCGs was more than 300bp in this study.

**Figure 1.**
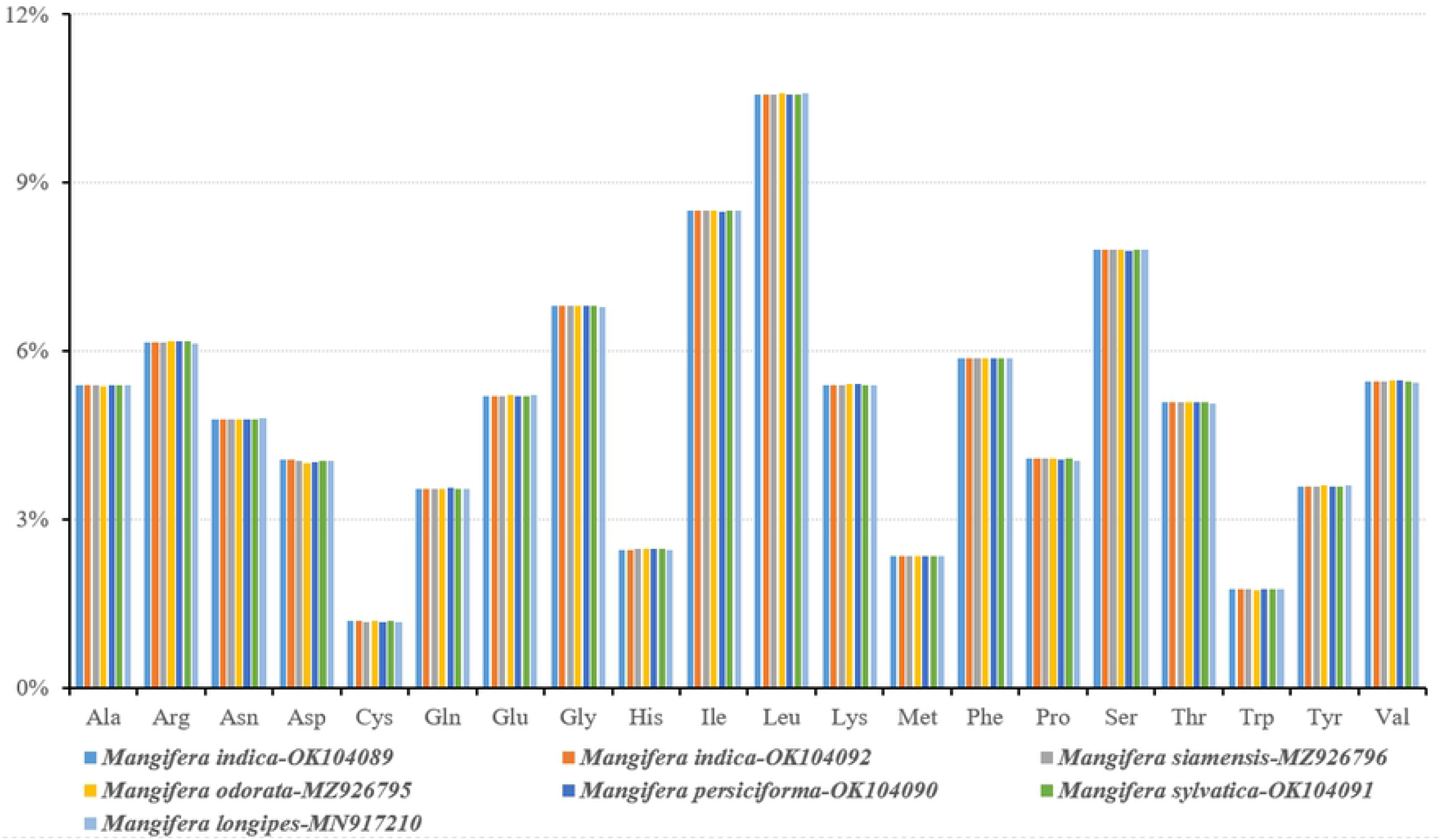
Amino acid proportion in 7 samples protein-coding sequences.

We detected all seven samples and found three results. Firstly, these seven samples could be divided into two groups by different amounts of editing sites. group1 included species ML210 and MP090. There were 61 RNA editing sites in 20 protein-coding genes in each species, with 44 occurring in the second nucleotide position and 17 in the first nucleotide position. Group 2 included other species; in this group, there were 60 RNA editing sites in 19 protein-coding genes in each species, with 45 occurring in the second nucleotide position and 15 occurring in the first nucleotide position. The different protein-coding gene between groups 1 and 2 was gene *ccsA*. Secondly, all RNA editing occurred through conversion of C to T and no other conversions. The third result was that gene *ndhB* had the most RNA editing sites in all species, and in these RNA editing sites, most of them occurred the conversions of serine to leucine (table S2).

We compared 35 protein-coding genes in 7 samples, then calculated the Ka/Ks and got over 3000 values, and we found that most of the Ka/Ks values were under 1. However, there still were 44 Ka/Ks values >1. When species MI089 was compared with others, there were five genes (*atp*B, *cem*A, *clp*P, *ndh*D and *pet*B) in 3 species (ML210, MO795 and MP090) had the Ka/Ks value over 1, and a similar result also happened in MI092, MS091 and MS796. When species MO795 was compared with others, there were four genes (*atp*B, *cem*A, *ndh*D and *pet*B) in 6 species (ML210, MS796, MI089, MS796, MS091 and MP090) had Ka/Ks values over 1. When species MP090 was compared with others, there were five genes (*atp*B, *cem*A, *clp*P, *ndh*D and *pet*B) in 6 species (ML210, MS796, MI089, MO795, MS091 and MI092) had the Ka/Ks value over 1. When species ML210 was compared with others, there were five genes (*atp*B, *cem*A, *clp*P, *ndh*D, *pet*D, *pet*B and *ycf*15) in 6 species (MP090, MS796, MI089, MO795, MS091 and MI092) had the Ka/Ks value over 1. The highest value of Ka/Ks was 3.9142, which happened on gene *pet*B in ML210 compared with MI089, MI092, MS796, MP090 and MS091. In addition, based on the data, there were seven genes that the Ka/Ks value >1, they happened on gene *atp*B, *cem*A, *clp*P, *ndh*D, *pet*D, *pet*B and *ycf*15 compared among seven samples. From the result, we could know that most genes would be under stabilize during evolution, but seven genes would be suffered from the positive selection after divergence. (Figure 2, Table S3)

**Figure 2.**
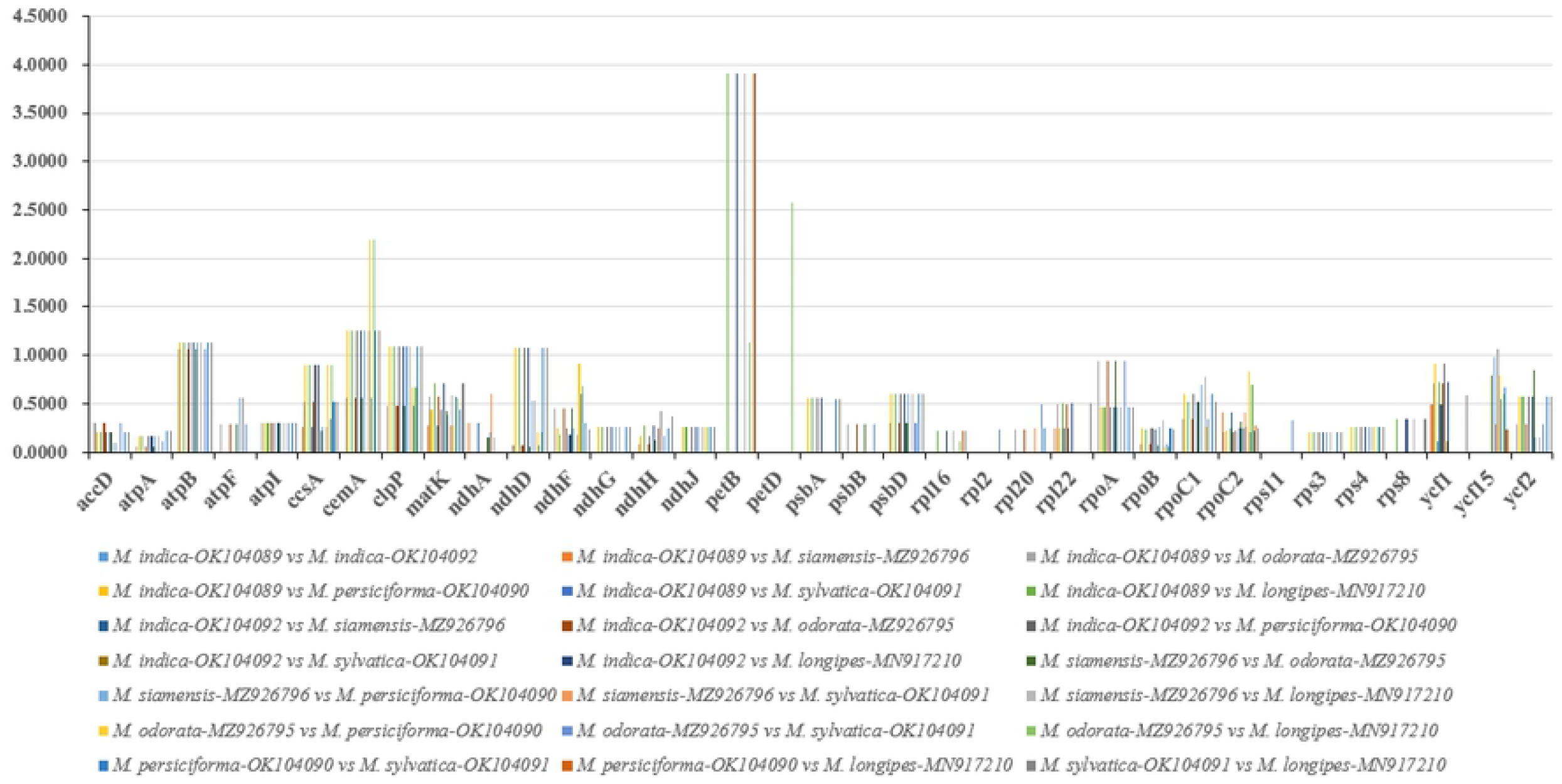
The value of Ka/Ks from a comparative of 7 samples.

### 3.3 Detection of Chloroplast Repeat Sequences and SSRs

This study identified two types of long repeats in 7 chloroplast genomes, which comprised forward and palindromic repeats. In contrast, no inverse repeats or complementary repeats were detected (Table S4), and the number of palindromic repeats was much more than forward repeats, covering more than 60%. Based on the result, the number of long repeats was around 10 to 20 pairs, MO795 had the least long repeat and ML710 had the longest repeats. Most of the repeats (93.8%) varied from 30 to 52 bp in length, and only a pair of repeats were more than 26360 bp in each species and were IR regions. Around these long repeats, more than 50% were located in LSC or SSC regions, and 7 chloroplast genomes were the same about this. The long repeats are mainly distributed around gene *rps*16, *rps*19, *ycf*1, *ycf*2, *ndh*A, *psa*A, *psa*B and *pet*D(table S4).

We analyzed seven samples separately and got different results for different species. 73 SSRs in MI089, 73 SSRs in MI092, 65 SSRs in MS796, 75 SSRs in MO795, 69 SSRs in MP090, 74 SSRs in MS091 and 66 SSRs in ML210. In these seven samples, the mononucleotides covered the most SSRs, and they covered around 60.87% - 67.57%. MS091 had the most mononucleotides, and it had 50 SSRs in mononucleotide type. The trinucleotides covered the second most SSRs, covering around 10.81%-16%. MO795 had the most trinucleotides, and it had 12 SSRs in trinucleotides type. The type of tetranucleotides covered the third most SSRs, round 10.81% - 15.94%. The dinucleotides covered around 4.35%-9.59% SSRs, and the type of hexanucleotides covered around 1.33%-1.54% SSRs. For the pentanucleotide type SSRs, we did not detect this type of SSRs in species MS796, MS091, ML210 and MO795. In MI089, MP090 and MI092, we detected one pentanucleotide type SSR, covering around 1.37-1.45% SSRs. Based on the result, we could find that MI089 and MI092 had the same amount SSRs even the quantity of the different types of SSR. In addition, C/G covered around 4.17-10.87% in different species in the mononucleotides. Most of the mononucleotides still were A/T. Furthermore, we also identified them in 10 PCGs and one tRNA gene; however, not all species had tRNA with SSR, MO795, ML210 and MP090 only contained 10 PCGs (totally, they distributed in different species.), but in others, there was a tRNA (gene *trn*K-UUU) containing SSRs (Table S5).

### 3.5 IR contraction and expansion

To further understand the structural characteristics of 7 chloroplast genomes, sequence alignments were performed using seven sequenced chloroplast genomes of *Mangifera* species, including ML210, MP090, MS796, MI089, MO795, MS091 and MI092.MO795 had the largest chloroplast genomes and LSC regions (158,889 and 87,708 bp, respectively). All five species have IR regions of similar sizes (26,360–26,389 bp) and LSC and SSC regions of varying sizes (Table 1). Variations in chloroplast sequence lengths among *Mangifera* species may be due to the different lengths of the LSC and SSC regions. Detailed comparative analysis of the junctions of the IRs and two single-copy regions, including LSC/IRA (JLA), LSC/IRB (JLB), SSC/IRA (JSA), and SSC/IRB (JSB), were conducted along with pla*cem*ent of adjacent genes in the chloroplast genomes of 7 *Mangifera* species. Seven genes, *rpl*22, *rps*19, *rpl*2, *ycf*1, *ndh*F, *trn*H and *psb*A, were detected at the junction of the LSC and IRs (Figure 3).

**Figure 3.**
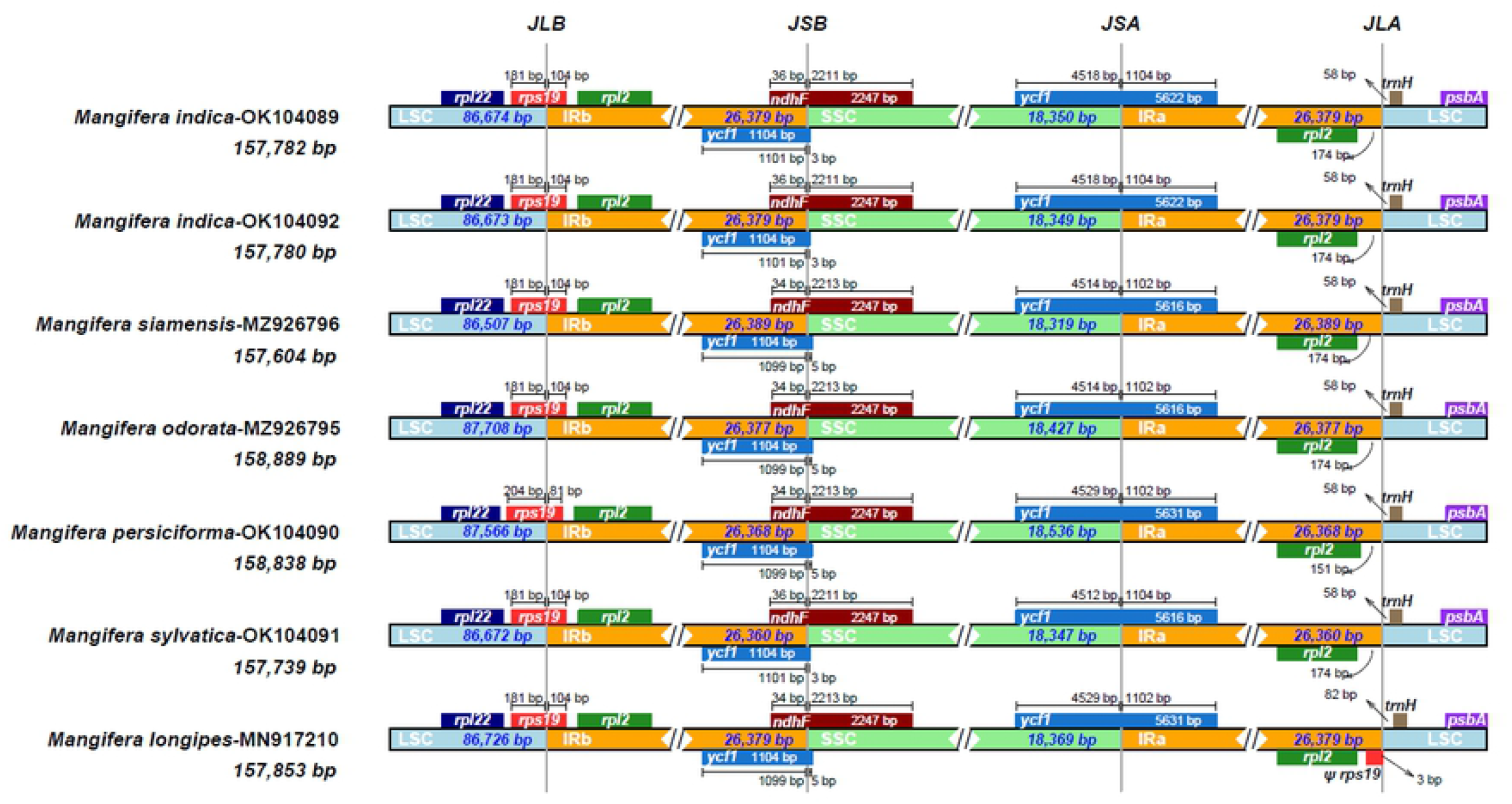
the four junctions of regions in chloroplast genomes of 7 *Mangifera* samples.

In MP090, *rps*19 is entirely located within the IR regions at 204 bp from JLB; however, in others, *rps*19 is closer (181 bp) to JLB within the IR regions. In MI089, MI092 and MS091, *ycf*1 is entirely located in IR regions 3 bp away from JSB and others were 5 bp away from JSB, however in MI089 and MI092, *ycf*1 was 4518 bp away from JSA, in MS796 and MO795, it was 4514 bp, in MS091, it was 4512 bp and others were 4529 bp. in ML210, *rps*19 was closer to JLA than *rpl*2, and the distance was 3 bp. However, in others, *rpl*2 was closer to JLA and the distance was around 151-174bp in different species. The gene *ndh*F spanned JSB in all species, but the distance to JSB was around 34-36 bp. And the location of *trn*H was on LSC in all species, and the distance to JLA was around 58-82 bp.

### 3.6 Sequence Divergence Analysis

We further analyzed the differences in the chloroplast sequences of 7 *Mangifera* species using mVISTA, with MI089 as a reference. The results indicated that the chloroplast genome sequences of *Mangifera* had very high sequence similarities, as shown in Figure 4. In addition, most differences happened in intergenic regions, but the coding regions were very calm. There were three different areas in the coding region; however, the differences were insignificant. They were the protein-coding genes *ycf*1 and *ycf*2, and these differences happened among species MS796, MO795, MP090, MS091 and ML210. However, significant differences were identified in some intergenic regions. There were three highly variable regions: *trnH-psb*A, *ycf4-ccmA* and *ndh*F-*rpl*32. (Figure 4).

**Figure 4.**
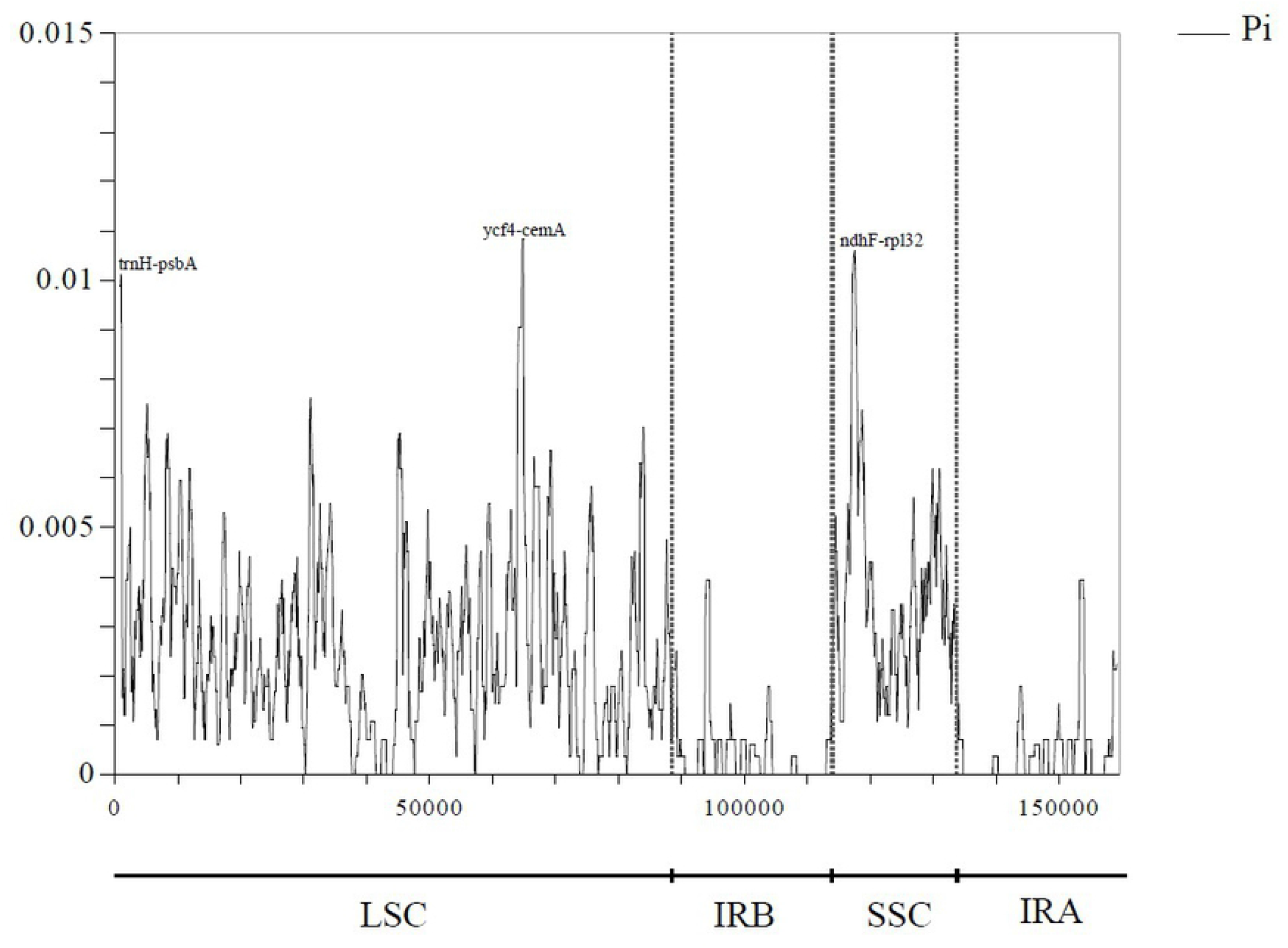
Global alignment of 7 chloroplast genomes of *Mangifera* using mVISTA.Y-axis indicates the range of identity (50–100%). Alignment was performed using MI089 as a reference.

We found that both the LSC and SSC regions were more divergent than the IR regions in the chloroplast genome of *Mangifera* species, with the three most variable regions (Pi > 0.010) were *trnH-psbA* and *ycf4-ccmA* and distributed in the of the LSC region, *ndh*F-*rpl*32 distribute in SSC regions. IR regions were no variable regions with Pi>0.005, and they were more stable and conservative than the other two regions. other six regions that showed higher variability (0.01 > Pi > 0.006) were distributed in the LSC region, including IGS *trn*K-UUU-*rps*16, *rps*16-*trn*Q-UUG, *pet*N-*psb*M, *trn*G-GCC-*psb*Z and *rpl*36-*rps*8. The high variability in these regions, especially in the LSC and SSC regions, provided information on diversity to develop markers for molecular classification and phylogenetic analysis of *Mangifera* species.

There were two same results in both Figure 4 and 5. They were the high divergent areas and high divergent regions. we could find that protein-coding genes *ycf*1 and *ycf*2, IGS *trn*H-*psb*A, *ycf*4-*ccm*A, *ndh*F-*rpl*32, *trn*K-UUU-*rps*16, *rps*16-*trn*Q-UUG, *pet*N-*psb*M, *trn*G-GCC-*psb*Z and *rpl*36-*rps*8 were mentioned in these section, especially IGS *trn*H-*psb*A, *ycf*4-*ccm*A, *ndh*F-*rpl*32 were mentioned in both figures. Furthermore, single copy regions were indicated that they were more divergent than others. In previous studies, many terms provided the evidence, such as the study of Dr. Fu, Dr. Trofimov’s team, Dr. Su’s team and Dr. Song’s team [41–44]. In their study, all the results of sequence divergent analysis had the same point: the most divergent areas distributed in single copy regions, either large single copy or small single copy. In addition, the high divergent areas in this study were covered by their studies, and the only difference was the different Pi values. The results of either previous studies or this study could state such opinions: the complete cp genomes in the same genus or families were very conservative, and the high divergent areas were in the limited range. By the way, in Dr. Machado’s study [41], they used this gene to analyze the phylogenetic relationship, and we also indicated that the gene *ycf*2 was more divergent than others in this study, so we tried to use this gene to verify the method mentioned in Dr. Machado’s study.

**Figure 5.**
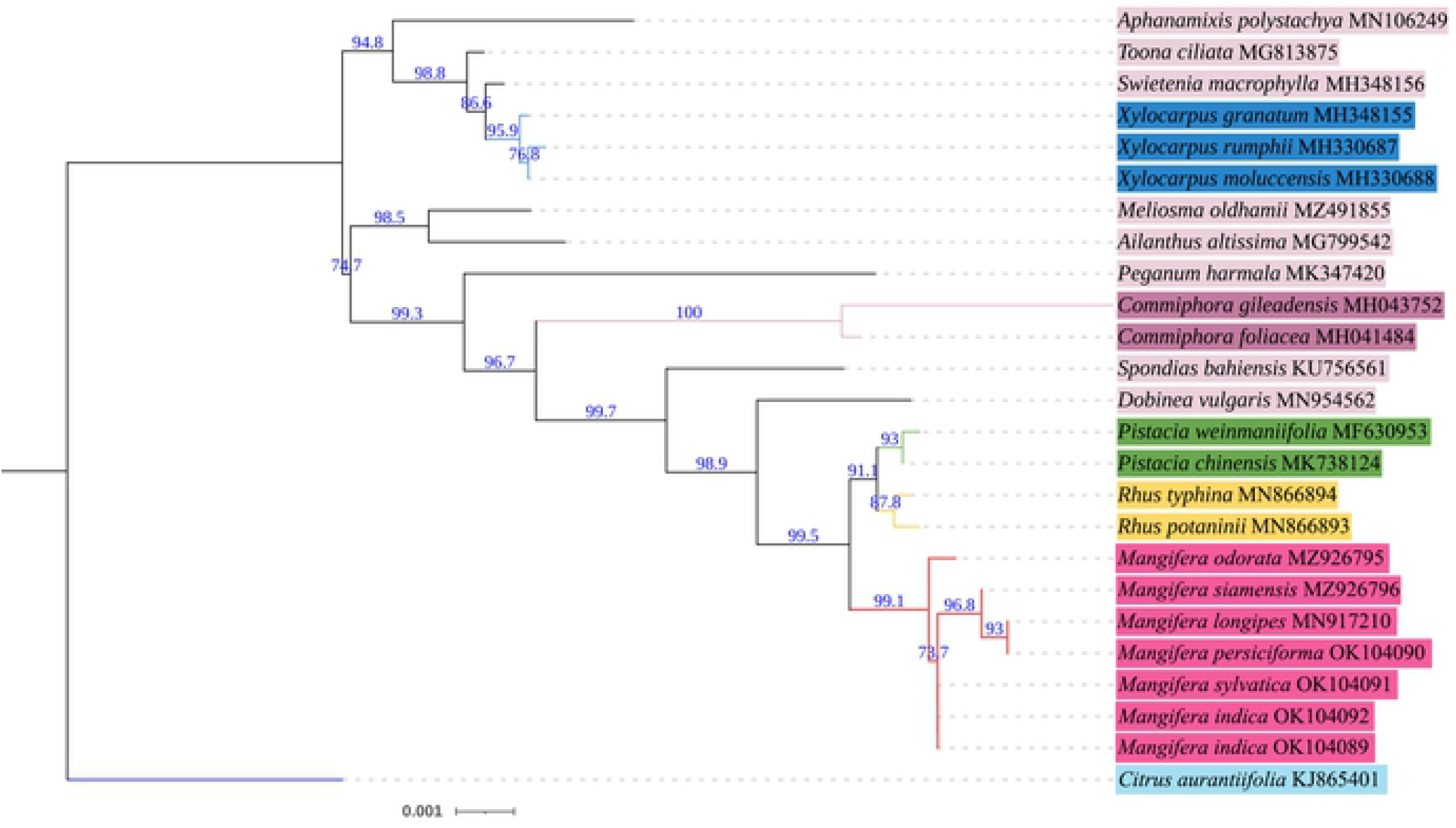
Nucleotide variability (Pi) values among chloroplast genomes of 7 *Mangifera* samples.

### 3.7 Phylogenetic Analysis

In this study, we constructed a phylogenetic tree and confirmed the relationship between 7 samples of *Mangifera* and 18 other angiosperm species with *Citrus aurantiifolia* (KJ865401) as an outgroup using the protein-coding gene *ycf*2 and complete chloroplast genome sequences separately (Figure 6 and Figure 7). We got similar results in two ways. In the result based on the gene *ycf*2, in 7 samples of *Mangifera*, ML210 and MP090 had a closer relationship than others; MS796 had a closer relationship with ML210 and MP090 than others. MS091, MI092 and MI089 had a closer relationship than others. And these six species also had a closer relationship than MO795. To compare the other 18 species, we could find that the genus *Mangifera* had a closer relationship with the genus Rhus and Pistacia. However, based on the complete chloroplast genome, there was a slight difference between MO795 and MS796 in Figure 7; MO795 was closer to MP090 and ML210 than MS796. Then we just used the 86 PCGs in each of these seven chloroplast genomes to construct a phylogenetic tree. In this tree, we did not use any outgroup, and we just used this phylogenetic tree to check the results from above. Because the difference between Figure 6 and Figure 7 is mainly in the genus *Mangifera*, the relationships of species between a genus or in their genus were the same. From 3 figures (Figure 6–8), the result based on 86 PCGs supported the result based on the complete chloroplast genome. We used the *ycf*2 gene to construct the phylogenetic tree and got a similar result between the genus. It meant that it could be possible to use gene *ycf*2 confirmed relationship when we could not obtain the complete chloroplast genome. In Dr. Machado’s study, they used the same method, which used the gene *ycf*2 in phylogenetic analysis, to construct the phylogenetic tree among the family Myrtaceae and got the same result to the result based on the complete cp genome and protein-coding genes [45]. So, this result could support Dr. Machado’s study at the family level. However, we could see some differences between these three methods. So at the genus level, *ycf*2 was not accurate enough compared with the methods based on the complete cp genome and proteincoding gene. We still need more work to verify the accuracy of the way base on gene *ycf*2. Because in this study, the result based on gene *ycf*2 could not be supported by other results.

**Figure 6.**
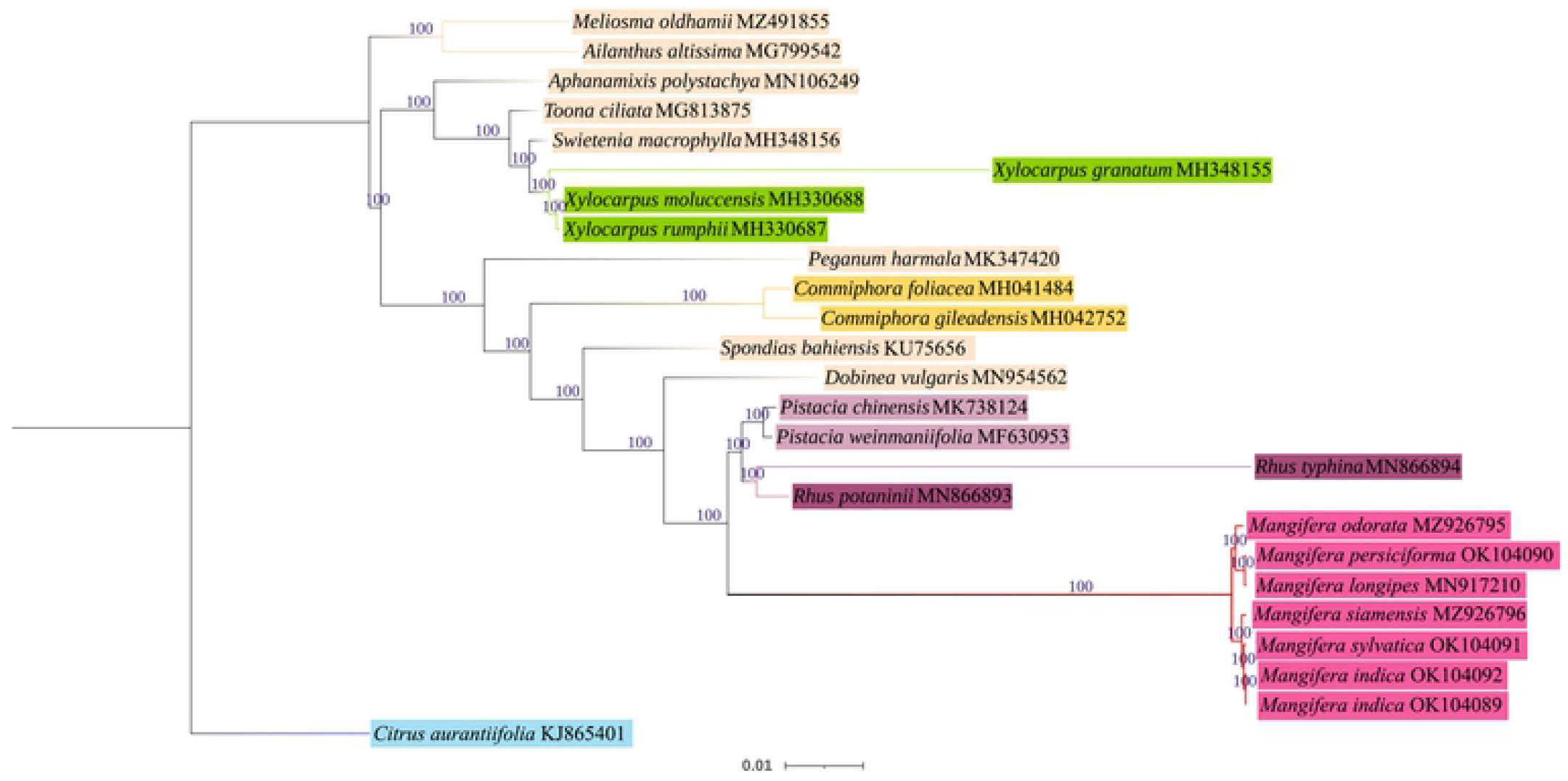
Phylogenetic tree inferred from maximum likelihood (ML) based on the gene *ycf*2 from 25 species.

**Figure 7.**
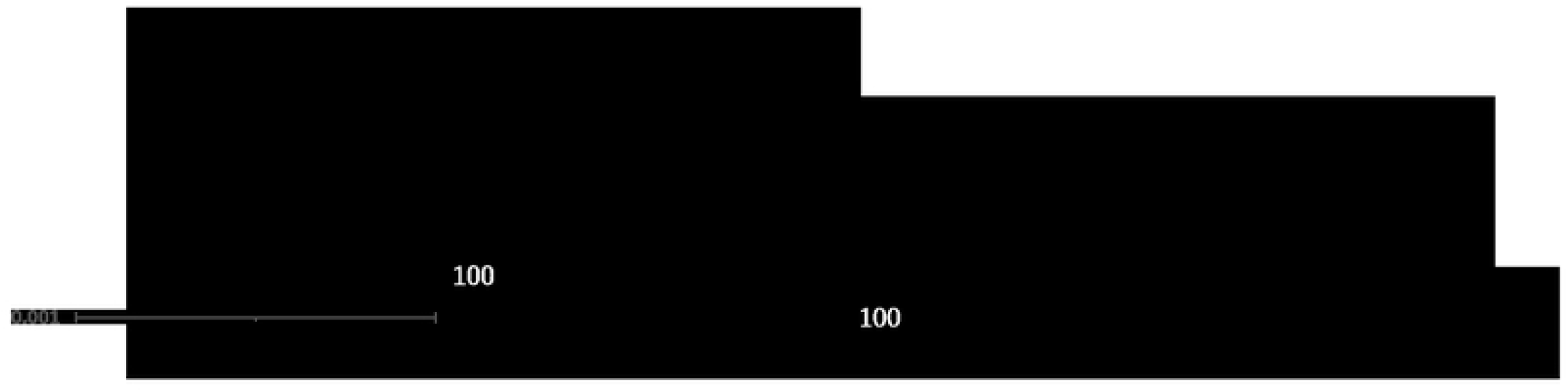
Phylogenetic tree inferred from maximum likelihood (ML) based on the 25 complete chloroplast genomes.

Phylogenetic tree inferred from maximum likelihood (ML) based on the 86 PCGs from complete chloroplast genomes of 7 *Mangifera* samples.

## 4 Conclusion

In this study, after sequenced and analyzed the species of *Mangifera*, we found that the size of the complete chloroplast of these seven samples was around 157,604 bp to 158,889 bp. The largest one was MO795, and the smallest one was MS796. Most of them contained 88 protein-coding genes, eight rRNA and 37 tRNA, except ML210, which only had 36 tRNA and 86 protein-coding genes. Through the RNA editing and Ka/Ks calculating, we found the species could be divided into two groups, and the difference between the two groups was the protein-coding gene *ccs*A. Moreover, all RNA editing occurred through conversion of C to T, and gene *ndhB* had the most RNA editing sites in all species. In Ka/Ks analysis, the gene *atp*B, *cem*A, *clp*P, *ndh*D, *pet*D, *pet*B and *ycf*15 would be suffered from the positive selection after divergence. We also find the IR regions in these seven samples were very conservation through IR contraction and expansion and Sequence Divergence Analysis. Finally, we tried to confirm the relationship between 7 samples of *Mangifera* in Angiosperms in 3 different ways. Then we got that ML210 and MP090 had a closer relationship than others, MS796 had a closer relationship with ML210 and MP090 than others. At the same time, when we did the phylogenetic analysis, we verified the accuracy of the method of phylogenetic analysis based on gene *ycf*2. We found it was not more accurate at the genus level than the method based on complete cp genome and protein-coding genes.

## Declarations

### Availability of data and materials

The chloroplast genome sequence was under the accession number: OK104089-OK104092, MZ926795-MZ926795 and MN917210 on the NCBI website. Supplementary materials can be found on the website.

Consent for publication: the details/images/videos will be freely available on the internet and may be seen by the general public

### Author Contributions

Yujuan Tang, Shixing Luo, Yu Zhang, Ying Zhao, Riwang Li, Limei Guo, Guodi Huang, Aiping Gao, Jianfeng Huang have equal Contributions

## Acknowledgments

We sincerely thank Total Genomics Solution Limited, SZHT. for performing the high throughput sequencing.

## Ethics approval and consent to participate

Not applicable

